# Reversion of pyrethroid resistant phenotypes in *Aphis glycines* by topical delivery of dsRNA targeting resistance alleles at the vgsc locus

**DOI:** 10.64898/2026.07.03.736413

**Authors:** Bruna Wojahn, Jonas André Arnemann, Matthew Elliott O’Neal

## Abstract

**BACKGROUND:** The soybean aphid, *Aphis glycines* Matsumura (Hemiptera: Aphididae), is a pest of soybean in North America that can cause significant yield loss when outbreaks are not managed. Current management tactics primarily rely on inexpensive pyrethroids, but the sustainability of this option is threatened by insecticide-resistance in *A. glycines* populations across the Upper-Midwest United States. Field-evolved resistance is associated with mutations in the voltage-gated sodium channel subunit h1 (*vgsc-h1*) gene.

**RESULTS:** Four double-stranded RNA (dsRNA) molecules, each matching the sequence of a *vgsc-h1* transcript variant (“Specific dsRNAs”), were topically applied to aphids with a genotype carrying the corresponding allele. The mortality of pyrethroid resistant aphids exposed to a Specific dsRNA increased in a dose-dependent manner when applied alone or with a constant concentration of lambda-cyhalothrin, plateauing at 1000 ng ul^-1^. Synergism was detected between two of four combinations of the Specific dsRNAs and lambda-cyhalothrin. These results were mirrored by the topical application of a single dsRNA with the consensus sequence of all *vgsc-h1* variants (“Combined dsRNA”). Mortality was consistently higher in aphids treated with either Specific dsRNA or the Combined dsRNA, alone or with lambda-cyhalothrin, compared to insecticide alone. The number of nymphs produced per female treated with the Specific or Combined dsRNA alone decreased significantly compared to untreated controls.

**CONCLUSION:** This study demonstrates that the topical application of dsRNAs targeting *vgsc-h1* increases the susceptibility and reduces the reproductive capacity of pyrethroid resistant soybean aphids, potentially providing a novel tool for the management of insecticide-resistant aphid populations.

## 1 INTRODUCTION

Soybean aphid, *Aphis glycines* Matsumura (Hemiptera: Aphididae), is an invasive pest of soybeans, *Glycine max* (Fabales: Fabaceae), in North America that was introduced from eastern Asia in the early 2000s.^1^ The soybean aphid lifecycle is holocyclic, alternating between asexual and sexual reproductive phases, and heteroecious, requiring an alternate host.^2^ Host switching involves overwintering on buckthorn, *Rhamnus cathartica* (Rosales: Rhamnaceae), followed by winged alates that migrate to *G. max* in the spring and summer, where parthenogenetic females produce approximately 15 asexual generations. Aphid populations can increase rapidly in the asexual generation.^3^ Aphids use a piercing-sucking stylet to feed on phloem through leaves, stems and pods which reduces plant vigor, and secretes honeydew that often leads to the growth of sooty mold that decreases plant photosynthetic capacity. This injury causes a decrease in plant height, pod set, and yield.^1^ Aphid outbreaks can reduce yield up to 40% if left unmanaged, though outbreaks typically reduce yield by approximately 13% in the state of Iowa.^4,5^

The intensive use of synthetic insecticides to manage *A. glycines* has contributed to the increased prevalence of resistance, resulting in failures of pyrethroids to manage outbreaks in the Upper Midwest region of the United States.^6–8^ Mutations in the voltage-gated sodium channel (*vgsc*) gene cause nonsynonymous (amino acid changing) substitutions in domain 2, segments 4 to 6 (D2S4-6) in the resulting protein, that alter pyrethroid binding and are associated with resistance to pyrethroids in several insect species.^9,10^ The pyrethroid knockdown resistant (*kdr*) phenotype in the housefly, *Musca domesticus* (Diptera: Muscidae), is associated with a leucine (L) to phenylalanine (F) change at position 1014 of the *vgsc* protein (L1014F).^11^ The positions of mutations in the *M. domesticus vgsc* protein are used as a standardized reference for orthologous changes. A greater level of resistance relative to the *kdr* phenotype, referred to as “super-knockdown resistant” (*skdr)*, is conferred by L1014F in conjunction with a secondary mutation at position 918,^11^ and exemplified in changes of the wildtype (*w*+) methionine (M) to leucine (L; M918L) or isoleucine (I; M918I) among insects.^12^ The *vgsc* protein in aphids is a heterodimeric complex composed of H1 and H2 subunits encoded by *vgsc-h1* and *-h2* genes, respectively.^13,14^ Mutations in the *vgsc*-*h1* D2S4-6 coding region are associated with pyrethroid-resistance among aphids (Hemiptera: Aphididae). For instance, the L1014F *kdr* mutation is associated with pyrethroid resistance in *Aphis gossypii*,^15^ *Myzus persicae*,^16^ and *Sitobion avenae*.^17^ Additionally, the M918I with L1014F, *skdr* genotype is found in *M. persicae*.^18^ A substitution causing a leucine to methionine change at position 925 (L925M) is associated with pyrethroid resistance in *Acyrthosiphon pisum*.^19^ In *A. glycines*, the M918I, M918L, L925M, and L1014F mutations are described in pyrethroid-resistant populations,^14,20,21^ wherein dominant mutations were mainly found in heterozygotes with exception of a homozygous *kdr* genotype. These nonsynonymous mutations are associated with reduced efficacy of pyrethroids against *A. glycines* in field populations.^20^

RNA interference (RNAi) is an endogenous gene regulatory mechanism found across Eukaryotes, including insects.^22,23^ This process modulates expression by either causing the degradation of specific messenger RNA (mRNA) or interfering with translational machinery of specific mRNA.^24–26^ Double-stranded RNA (dsRNA) molecules induce the RNAi response,^27^ leading to sequence-specific silencing of corresponding mRNAs. This discovery was developed into a novel reverse-genetic tool to repress gene function and study subsequent phenotypes.^22^ RNAi-induced knockdown of genes critical for survival demonstrates the creation of novel insecticides,^28–30^ that could selectively target pest species while preserving the health of non-target organisms.^31^ The use of RNAi technology for the creation of novel insecticides has the potential to enhance the precision and effectiveness of pest control strategies.^32^ The first commercially available RNA-based insecticide was developed to silence the *snf*7 gene of western corn rootworm, *Diabrotica virgifera virgifera* (Coleoptera: Chyrsomelidae) and was deployed through genetically-engineered maize hybrids.^33,34^ A foliar-applied insecticidal RNA product named Calantha is marketed for management of the Colorado potato beetle, *Leptinotarsa decemlineata* (Coleoptera: Chrysomelidae).^35,36^

The prevalence of pyrethroid resistance in soybean aphid populations threatens the long-term utility of this management option for farmers across a major soybean producing region in the Midwest United States. Novel strategies that preserve or extend the efficacy of pyrethroids could help protect profit margins, as this active ingredient is one of the least expensive insecticides available to farmers in the US. Our prior work demonstrated that the topical application of dsRNAs targeting transcripts of the *vgsc-h1* reduces intracellular transcript quantities (knockdown) and increases mortality among pyrethroid resistant aphids.^37^ The current study integrates these dsRNAs into an insecticidal framework. This is accomplished by determining what dose of a topically applied dsRNA optimizes the performance among different dsRNA constructs that are either 1) a “Specific dsRNA”, each with 100% sequence identity to one of three different field-evolved *A. glycines vgsc-h1* allele variants associated with pyrethroid resistance, or 2) a “Combined dsRNA” with a consensus sequence of all variant alleles. Insecticidal properties of these novel RNAs alone or with a pyrethroid serve as a foundation for a field-ready strategy to manage pyrethroid resistant aphid populations when used within an integrated pest management strategy.

## 2 MATERIALS AND METHODS

### 2.1 Isofemale lines

Collection and identification of the genotypes and phenotypes of five parthenogenetic (clonal) *A. glycines* isofemale lines used in this study were described previously (Table 1).^14,20^ In brief, each isofemale line was initiated from a single, field-collected female. Laboratory assays were previously conducted with each line to determine the resistance ratio (RR) to pyrethroids with respect to the susceptible isoline, SBA-Boone-2019-ISO (Table 1). For this study, the isolines were numbered in order of increasing RR, with isoline I being susceptible, and isolines II through V having a successively greater estimated RR. Isoline I and V were initiated with aphids collected at fields near Boone, IA and Darwin, MN, USA, respectively, prior to any pyrethroid application during that growing season. Isolines II, III and IV were each started from a single female collected from different fields of *G. max* following a field-applied rate of lambda-cyhalothrin (Warrior II, Syngenta Crop Protection, Greensboro, NC). All isolines were propagated in the laboratory on the *G. max* cultivar NK S24-K2 (Syngenta Crop Protection) in a Percival growth chamber (Percival Scientific, Perry, Iowa, USA) at 25°C, 50% RH and 16:8 L:D. Colonies were maintained without exposure to insecticides until used in this study.

**Table 1.**
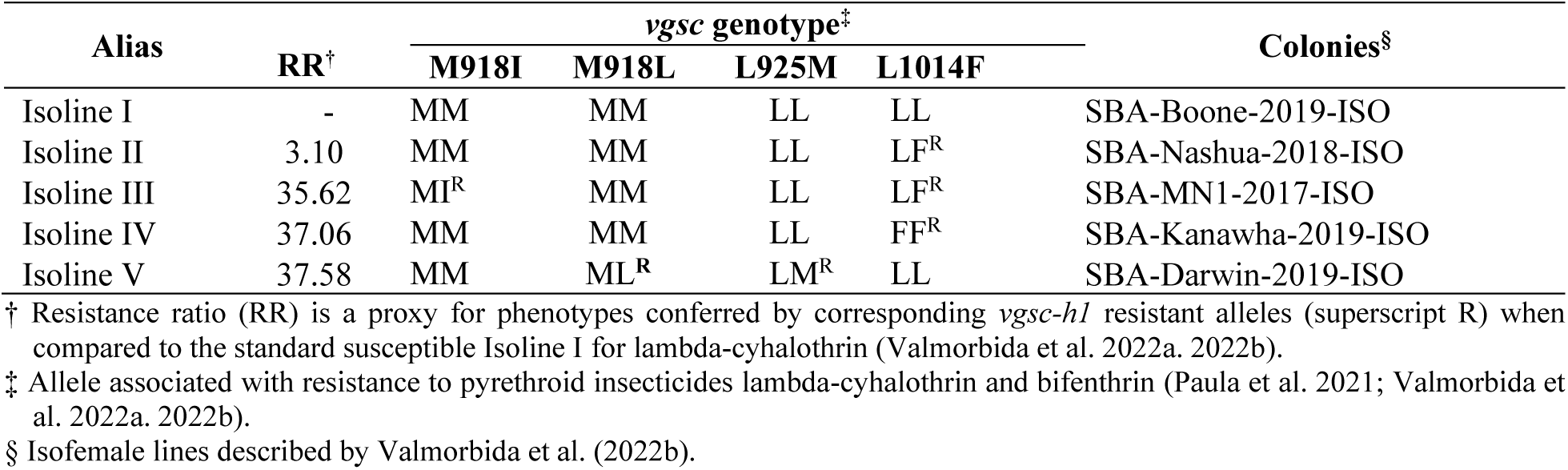
Isofemale lines of *Aphis glycines,* with varying genotypes related to the voltage-gated sodium channel subunit h1 (*vgsc-h1*) and lambda-cyhalothrin resistance level.

### 2.2 Site-directed mutagenesis

Individual dsRNA constructs were developed via site-directed mutagenesis of complementary DNA (cDNA) from native allele (transcript) sequence to include ≥ 1 nucleic acid change serving as template for subsequent production of *vgsc-h1* dsRNAs (ds*vgsc-h1*). These site-directed changes corresponded to those at positions 2782, 2784, 2803, and 3070 of the *A. glycines vgsc-h1* CDS (transcript model AG6007485-RA; Figure S1)^38^ that are predicted to cause amino acid changes M918L, M918I, L925M, and L1014F, respectively. All four mutations are associated with pyrethroid resistance in *A. glycines*.^21,14,20^ Transcripts used for site-directed mutagenesis were from Isoline I, which is homozygous for the *w*+ *vgsc-h1* allele, and Isoline IV, which is homozygous for the L1014F mutation. The cDNA from Isoline I was altered to match specific alleles that encoded 918I + 1014F (ds*vgsc-h1*^918I+1014F^) or 918L + 925M mutations (ds*vgsc-h1*^918L+^ ^925M^) (Figure S1). Isoline IV cDNA was used without modification as a template for a dsRNA with the 1014F mutation (ds*vgsc-h1*^1014F^). Isoline IV cDNA was also modified to combine all substitutions responsible for 918I, 918L, 925M, and 1014F amino acid mutations (Combined dsRNA; ds*vgsc-h1*^All^). These *dsvgsc-h1* sequences are also described in United States Patent US2024/059991.^37^

Total RNA was extracted from separate pools of 40 mixed age apterous aphids from Isolines I and IV submerged while alive in Trizol Reagent (Invitrogen, Waltham, MA, USA) using Direct-zol RNA Miniprep Kit (Zymo Research, Irvine, CA, USA) according to the manufacturer’s instructions. Purified RNA was then quantified by absorbance at 230 nm on a DeNovix DS-11 spectrophotometer (DeNovix Inc, Wilmington, DE, USA) and ∼0.5 to 1.0 µg used for first-strand cDNA synthesis primed by random hexamers using the GoScript^TM^ Reverse Transcription (RT) System (Promega, Madison, WI, USA) according to the manufacturer’s instructions. The resulting cDNA from each Isoline was used as template in separate reactions that amplified a 485 bp fragment that spanned positions 2711 to 3195 of the *A. glycines vgsc-h1* CDS in AG6007485-RA.^38^ These amplicons were generated in a 40.0 μl RT-PCR reaction volume with 1X LongAmp polymerase buffer (New England Biolabs, Ipswich, MA, USA), 1.5 mM MgCl_2,_ 60 μM dNTPs, 8.0 µl of a 1:100 dilution of first strand cDNA with nuclease-free water, 1.3 U LongAmp DNA polymerase (New England Biolabs), and 0.25 µM of each primer pair Ag6007485_2711t7-S and _3195t7-A (Table S1a) that each incorporated 3’ overhangs corresponding to the T7 promoter sequence (5’-TAA TAC GAC TCA CTA TAG GGA GA-3’). Reactions were run in a touchdown thermocycler program as described previously.^39^ A 5.0 µl aliquot of each reaction product was separated by 2% agarose gel electrophoresis to assess quality. The remaining product volumes were treated with Exonuclease I (ExoI) and Shrimp Alkaline Phosphatase (SAP) (New England Biolabs, Ipswich, MA USA) to remove residual primer and nucleotides as described previously^14^ and diluted 1:10^-7^ in nuclease-free water.

Site-directed mutagenesis was performed on diluted RT-PCR products from Isoline I via primer extension reactions using oligonucleotides AgΔ918I+Δ925M, or from Isoline IV using AgΔ918I or AgΔ918IL+Δ925M (Table S1). Primer extension reactions consisted of 1X LongAmp Reaction Buffer (New England Biolabs), 60 μM dNTPs, 0.3 µM of oligonucleotide AgΔ918I, AgΔ918I+Δ925M, or AgΔ918IL+Δ925M, 4.0 µl of the corresponding diluted RT-PCR amplicon, and 1.0 U LongAmp DNA polymerase (New England Biolabs) in a 40.0 μl volume. Reactions were incubated at 95°C for 1 m, then 30 cycles of 80°C for 30 s and 68°C for 1 m, followed by 68°C for 3 m. A 0.5 µl aliquot of each primer extension product was then PCR amplified using the primer pair Ag6007485_2711t7-S and _3195t7-A (TableS1) as described above, then 5.0 µl of these PCR products were separated by agarose gel electrophoresis as described above. A total of 5.0 µl from the remaining PCR product volume was treated with ExoI and SAP (New England Biolabs) as described above, diluted 1:20 with nuclease free water, and submitted to the Iowa State University DNA Facility (Ames, IA) for bi-directional sequencing on ABI3700 (Applied BioSystems, Waltham, MA USA) using primers Ag6007485_2711t7-S and _3195t7-A in separate sequencing reactions. Raw Sanger read data was analyzed as described previously^14,20^ and used to validate fixation of nucleotides by site directed mutagenesis. The PCR products with validated DNA sequence were used as templates in subsequent dsRNA synthesis reactions.

### 2.3 Double-stranded RNA synthesis

The PCR products with validated sequence were mixed with a 1.8X volume of Agencourt AMPure-XP Beads (Beckman-Coulter, Hercules, CA, USA) and purified according to the manufacturer’s protocol, then quantified using Quant-iT™ DNA Broad Range (BR) Assay Kits (Thermo Fisher Scientific, Waltham, MA, USA) with fluorescence measured on a DeNovix DS-11 (DeNovix Inc). Between 0.8 and 1.25 µg of purified template were then used in separate dsRNA synthesis reactions using the Invitrogen MEGAscript^TM^ T7 Transcription Kit (Thermo Fisher Scientific) according to manufacturer instructions. Transcription reactions were incubated at 37°C for 16 hrs. The subsequent synthesized dsRNA was purified using the Invitrogen MEGAclear^TM^ Transcription Clean-Up Kit (Thermo Fisher Scientific) and quantified using Quant-iT™ RNA Broad Range (BR) Assay Kits (Thermo Fisher Scientific) as instructed by the manufacturers.

### 2.4 dsRNA dose-response bioassay

The impact of varying concentrations of the ds*vgsc-h1*^1014F^ on mortality was evaluated among aphids from Isoline IV homozygous for the corresponding L1014F mutation, to determine the “optimal” concentration at which the dsRNA induces mortality when applied alone and with a constant concentration of lambda-cyhalothrin. For this, adults from Isoline IV were exposed to insecticide treatments via leaf-dip bioassays as described previously,^14^ following established guidelines to determine the response to insecticides.^40^ In brief, aphids were transferred to 3.8 cm diameter of a detached leaf disk from the *G. max* cultivar NK S24-K2 (Syngenta) dipped for 10 s into a 1.23 parts per million (ppm) solution of lambda-cyhalothrin in distilled water containing 0.05% (v/v) Triton X-100 (Alfa Aesar, Tewksbury, USA), diluted from a 0.5 mg ml^-1^ stock solution of lambda-cyhalothrin created in acetone. The constant 1.23 ppm corresponded to the lethal concentration sufficient to cause 90% mortality (LC90) in the *w*+ susceptible Isoline I.^14^ Leaf disks for treatments without insecticide and controls were immersed in a solution of 0.05% (v/v) Triton X-100 (Alfa Aesar).

Prepared leaf disks were then placed onto a bed of 1% w/v agar (Bacto™ Agar, Becton, Dickinson and Company, Franklin Lakes, USA) poured at the bottom of plastic souffle cups (29.6 ml; Choice Paper Company, New York, USA). Each cup was filled with 20 ml of agar, and a drop of distilled water was added to the top of the agar to increase the adherence of the leaf. Subsequently, 20 uninjured adult aphids were collected from Isoline IV colony (Table 1) and transferred to each leaf disk. The purified ds*vgsc*-*h1*^1014F^ was diluted to 0, 10, 100, 200, 400, 500, 600, 1000 and 2000 ng µl^-1^ in 1.0 X Tris-EDTA buffer pH 7.6 containing 0.05 % (v/v) Tween 20 (Tween® 20, Sigma-Aldrich, Saint Louis, MO, USA). A 0.2 μl droplet of ds*vgsc*-*h1*^1014F^ preparations was applied directly to the dorsal side of the abdomen on each aphid using a repeating dispenser (Hamilton PB600-1, Hamilton Co., Reno, NV, USA). Each cup with an infested leaf disk was considered an experimental unit to which treatments of the ds*vgsc*-*h1*^1014F^ alone or with lambda-cyhalothrin were applied. Treatments were assigned based on a randomized complete block design, with three replicates (cups) per treatment. One replicate of each treatment was conducted per day, with experimental date serving as the blocking factor. All cups were sealed with a ventilated lid and placed in a Percival growth chamber (Percival Scientific, Perry, Iowa, USA) at 25°C, 50% RH and 16:8 L:D, and mortality was measured at 6, 24, and 48 hours post-treatment. Mortality was estimated at 6 and 24 h by counting dead aphids based on their upright appearance without disturbance. At 48 h, aphids were turned on their backs and considered dead if unable to right themselves within 10 seconds. Additionally, the number of nymphs birthed by treated females was counted in each cup at 48 h and divided by the number of live adults at 24 h to estimate the average number of nymphs produced per adult.

### 2.5 Effects induced by specific and combined dsRNAs

Based on outcomes from the above ds*vgsc-h1*^1014F^ dose-response assays on Isoline IV, we elected to use a concentration of 1000 ng ul^-1^ to determine the impact of other Specific dsRNAs and the Combined dsRNA on aphid mortality when applied in the presence and absence of 1.23 ppm lambda-cyhalothrin. Leaf disks with a constant 1.23 ppm of lambda-cyhalothrin were prepared and infested with aphids from a single Isoline as described above. Aphids were treated with a 0.2 μl droplet of a dsRNA having a sequence that matched the resistant allele of its corresponding genotype (Specific dsRNA; Table 2), as well as the ds*vgsc*-*h1*^All^ that combined all nucleotide changes (Combined dsRNA; Figure 1). The allele-specific dsRNA treatments included: the ds*vgsc*-*h1*^1014F^ on *A. glycines* from Isoline II and IV that are heterozygous and homozygous for the L1014F mutation, respectively; the ds*vgsc*-*h1*^918I+1014F^ on Isoline III which is heterozygous for the M918I and L1014F mutations; and ds*vgsc*-*h1*^918L+925M^ on aphids heterozygous for M918L and L925M mutations in Isoline V. The ds*vgsc*-*h1*^All^ was applied to *A. glycines* from isolines I, II, III, IV and IV (Table 2). Aphid mortality and the number of nymphs were measured as described above across all treatments.

**Figure 1.**
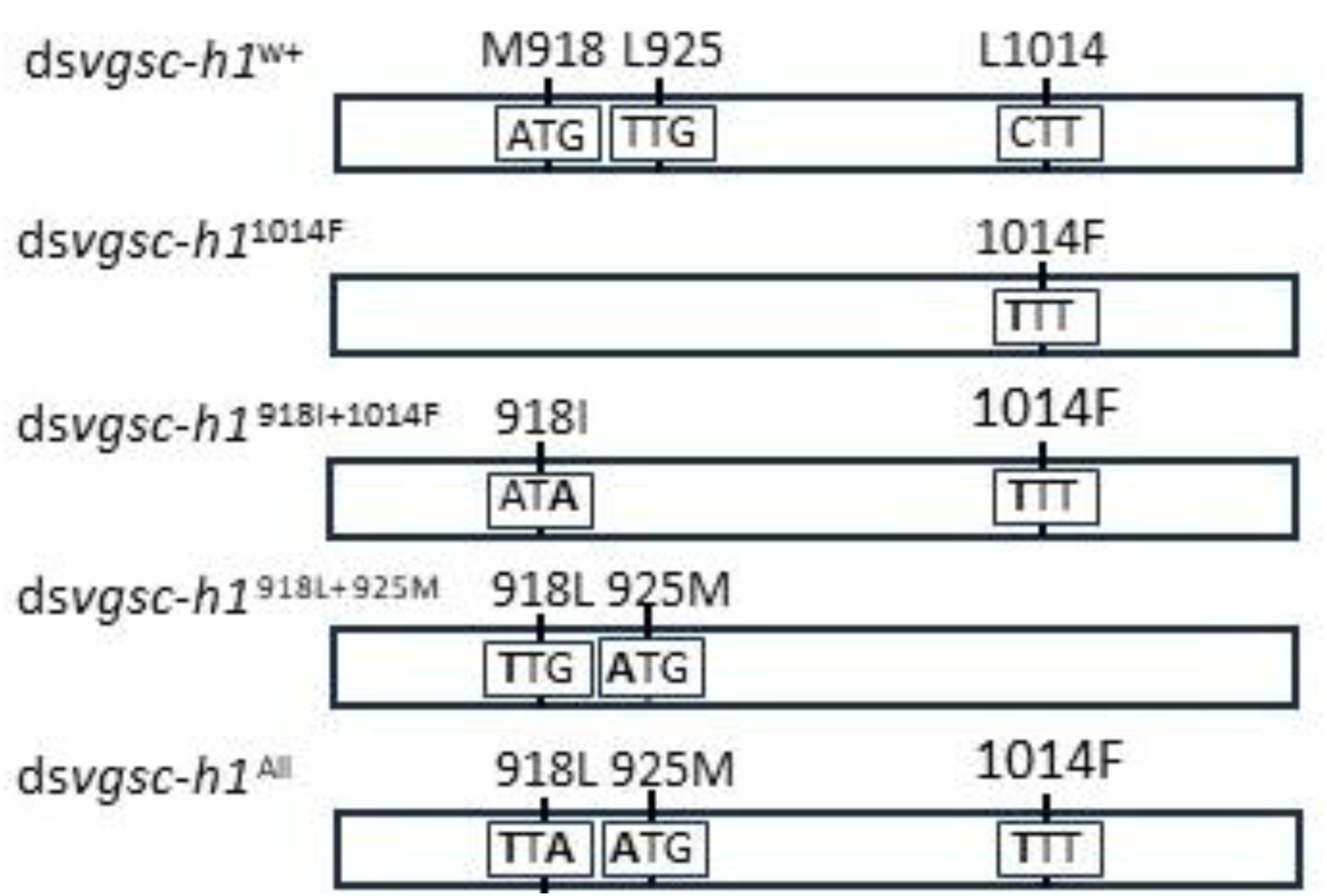
Diagram of double-stranded RNA (dsRNA) molecules that target alleles of the *Aphis glycines voltage-gated sodium channel* subunit h1 gene (ds*vgsc-h1*). Sequence of ds*vgsc-h1^w+^* matches that of the wildtype (*w*+) allele homozygous in Isoline I. Nucleotides that change among the ds*vgsc-h1*^1014F^, ds*vgsc-h1*^918I+1014F^, ds*vgsc-h1*^918L+925M^, and ds*vgsc*-*h1*^All^ with respect to ds*vgsc-h1^w+^*are in bold, and nucleotides enclose within boxes representing codon triplets (Remainder of dsRNA sequence remain unchanged from those at positions 2633 to 3195 of the transcript model AG6007485-RA; **Figure S1**). Predicted amino acids encoded by corresponding native *vgsc-h1* alleles associated with pyrethroid resistant are represented with respect to orthologous positions in the *vgsc* protein of *Musca domestica*.^59^ The ds*vgsc*-*h1*^All^ is an artificial construct that combines all four nucleotide substitutions found across *vgsc-h1* alleles associated with pyrethroid resistance.

**Table 2.**
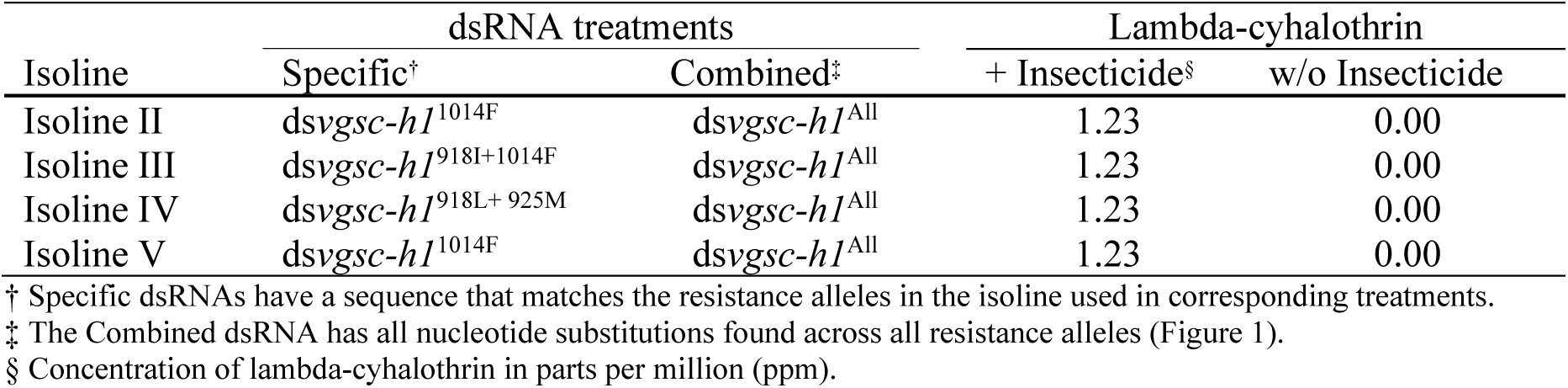
Double-stranded RNA (dsRNA) treatments applied to *Aphis glycines* isolines II, III, IV and V (Table 1).

### 2.6 Statistical analysis

Mortality was corrected using Abbott’s formula: [(X - Y)/ X] × 100 = percent control; where X = percent of living insects in the control and Y = percent of living insects in the treatment.^41^ Statistical analyses were performed using R software version 4.4.2 (R Foundation for Statistical Computing, Vienna, Austria) and Excel (Microsoft, Redmond, WA USA). Data were analyzed using generalized linear models (GLMs) with a factorial design including treatment effects and their interactions. A quasibinomial distribution with a logit link was used for mortality data, and a quasipoisson distribution with a log link was used for the number of nymphs per adult. When interactions were significant, post-hoc pairwise comparisons derived from the GLMs were performed. Comparisons were performed to determine the impact of the insecticide at each dsRNA concentration in the corresponding dose-response bioassay, as well as differences between Specific dsRNAs (ds*vgsc-h1*^918L+925M^, ds*vgsc-h1*^918I+1014F^, or ds*vgsc-h1*^1014F^) and Combined dsRNA (ds*vgsc-h1*^All^) treatments within an isoline, across treatments with and without lambda-cyhalothrin. P-values were adjusted for multiple comparisons using the Šidák method,^42^ and a significance level of 0.05 was used for all analyses.

## 3 RESULTS

### 3.1 Dose response bioassay

The mortality among *A. glycines* from Isoline IV increased significantly in a dose dependent manner when exposed to increasing concentrations of ds*vgsc*-*h1*^1014F^. We observed a significant interaction between dsRNA and lambda-cyhalothrin using GLM (p < 0.0001; Figure 2A), but no significant effect was detected for the insecticide alone. This latter result is expected as Isoline IV is resistant to lambda-cyhalothrin. Results of post-hoc pairwise Šidák comparisons showed that lambda-cyhalothrin at a constant 1.23 ppm dose had a positive effect on mortality when combined with ds*vgsc*-*h1*^1014F^ at concentrations ≥ 400 ng μl^-1^ (p < 0.0001; Figure 2A). The highest estimated mortality at 48 h post-application was 96% observed at 2000 ng ul^-1^ of ds*vgsc*-*h1*^1014F^ with lambda-cyhalothrin, while the lambda-cyhalothrin alone treatment (0 ng ul^-1^ of dsRNA + Insecticide) caused 11% mortality. The estimated 87% mortality at the 1000 ng μl^-1^ concentration of ds*vgsc*-*h1*^1014F^ + Insecticide was not significantly different from that at 2000 ng μl^-1^ + Insecticide. Based on these results, we elected to use 1000 ng μl^-1^ as the “optimal” concentration of dsRNA in all subsequent bioassays with allele-specific and Combined dsRNAs (200 ng effective dose in each 0.2 μl topically applied droplet).

**Figure 2.**
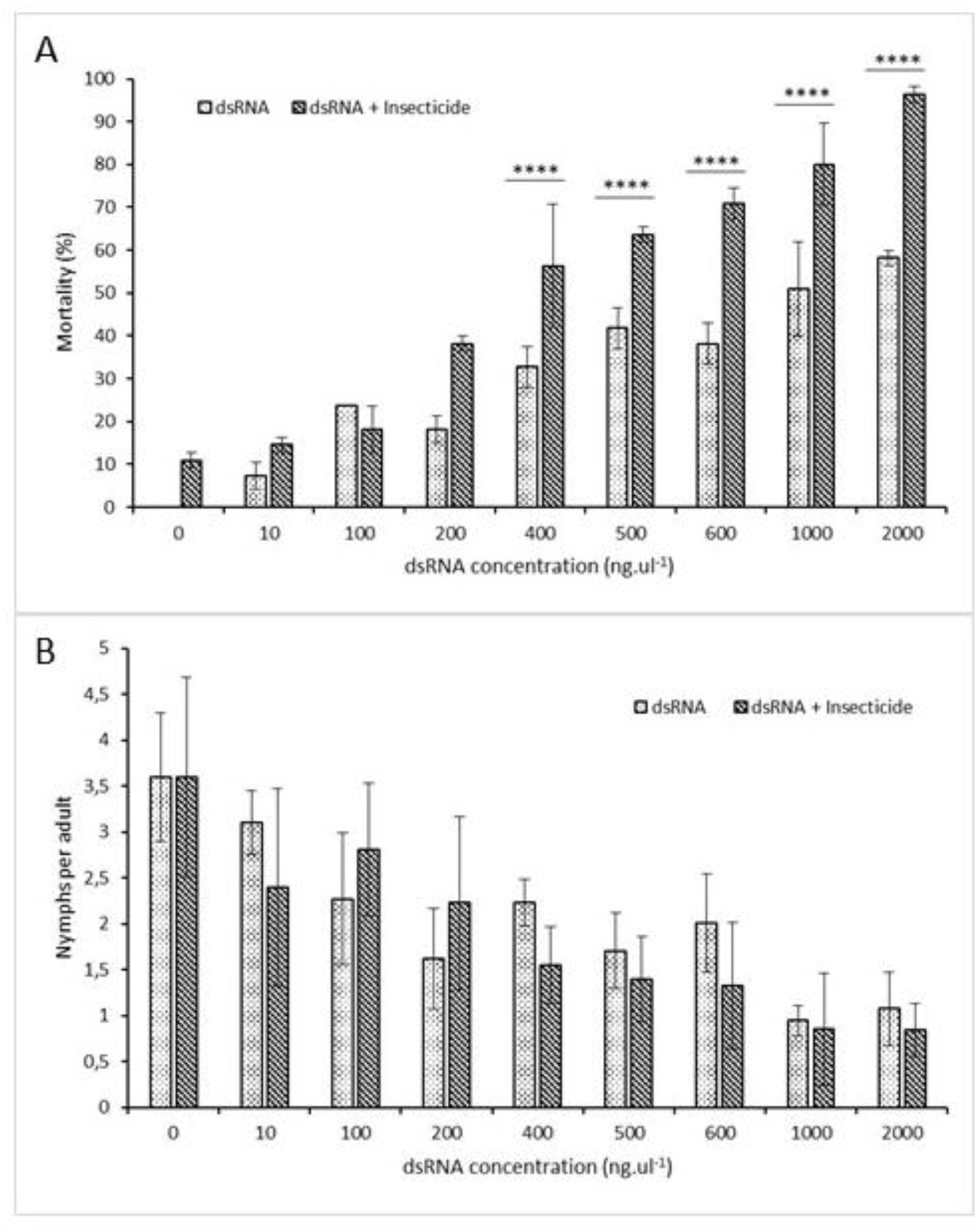
Dose-responses in mortality (A) and number of nymphs produced per adult (B) induced in *Aphis glycines* isoline IV, homozygous for the L1014F mutation (Table 1). Aphids were exposed to varying concentrations of ds*vgsc-h1*^1014F^. Each concentration of ds*vgsc-h1*^1014F^ was applied alone (dsRNA) or along with lambda-cyhalothrin at a constant concentration of 1.23 ppm (dsRNA + Insecticide). For mortality (A), a significant dose-dependent effect of dsRNA was detected (p < 0.0001). For reproduction (B), the effect of dsRNA concentration on the number of nymphs produced per adult was also significant (p = 0.0048). Asterisks indicate statistically significant differences between dsRNA and dsRNA + Insecticide treatments within the same concentration of dsRNA (p < 0.0001).

The number of *A. glycines* increased during the experiments due to the birth of nymphs from parthenogenic females, but this increase was suppressed among females receiving ds*vgsc*-*h1*^1014F^ in a dose-dependent manner (p = 0.0048; Figure 2B). In contrast, the control treatments with no insecticide and dsRNA reached an average of 90 aphids per experimental unit after 48 h, corresponding to an average of 3.6 nymphs produced per infested adult. Treatment with the ds*vgsc*-*h1*^1014F^ alone had a significant negative effect on reproduction. Reproduction was less than 1.0 nymph per adult at 48 h post-application for the 1000 and 2000 ng ul^-1^ ds*vgsc*-*h1*^1014F^ concentrations combined with the insecticide. We did not observe an interaction of the dsRNA with the insecticide.

### 3.2 Effects induced by specific and combined dsRNA sequences

Aphid mortality varied by dsRNA sequence and isoline (Figure 3A-D). The lowest mortality was consistently observed when the insecticide was applied alone. The insecticide alone caused 12% mortality in Isoline II, which has the lowest RR (3.1; Table 1) and did not exceed 5% mortality in the other resistant Isolines (III, IV and V) that each have a ten-fold greater RR than Isoline II. Specific and Combined dsRNAs alone produced 50 to 80% mortality among the resistant isolines, with no significant differences between dsRNAs within isoline. Mortality induced by treatments with the Specific and Combined dsRNA with lambda-cyhalothrin (+ Insecticide) was significantly greater compared to treatments with these dsRNAs alone in Isoline II (Figure 3A). Treatment of aphids from Isoline IV with Specific dsRNA + Insecticide produced significantly greater mortalities compared to either the Specific or Combined dsRNA alone, but not the Combined dsRNA + Insecticide (Figure 3C). In contrast, treatments with Specific or Combined dsRNA, alone or with the insecticide, were not significantly different from one another in Isoline III (Figure 3B) or V (Figure 3D).

**Figure 3.**
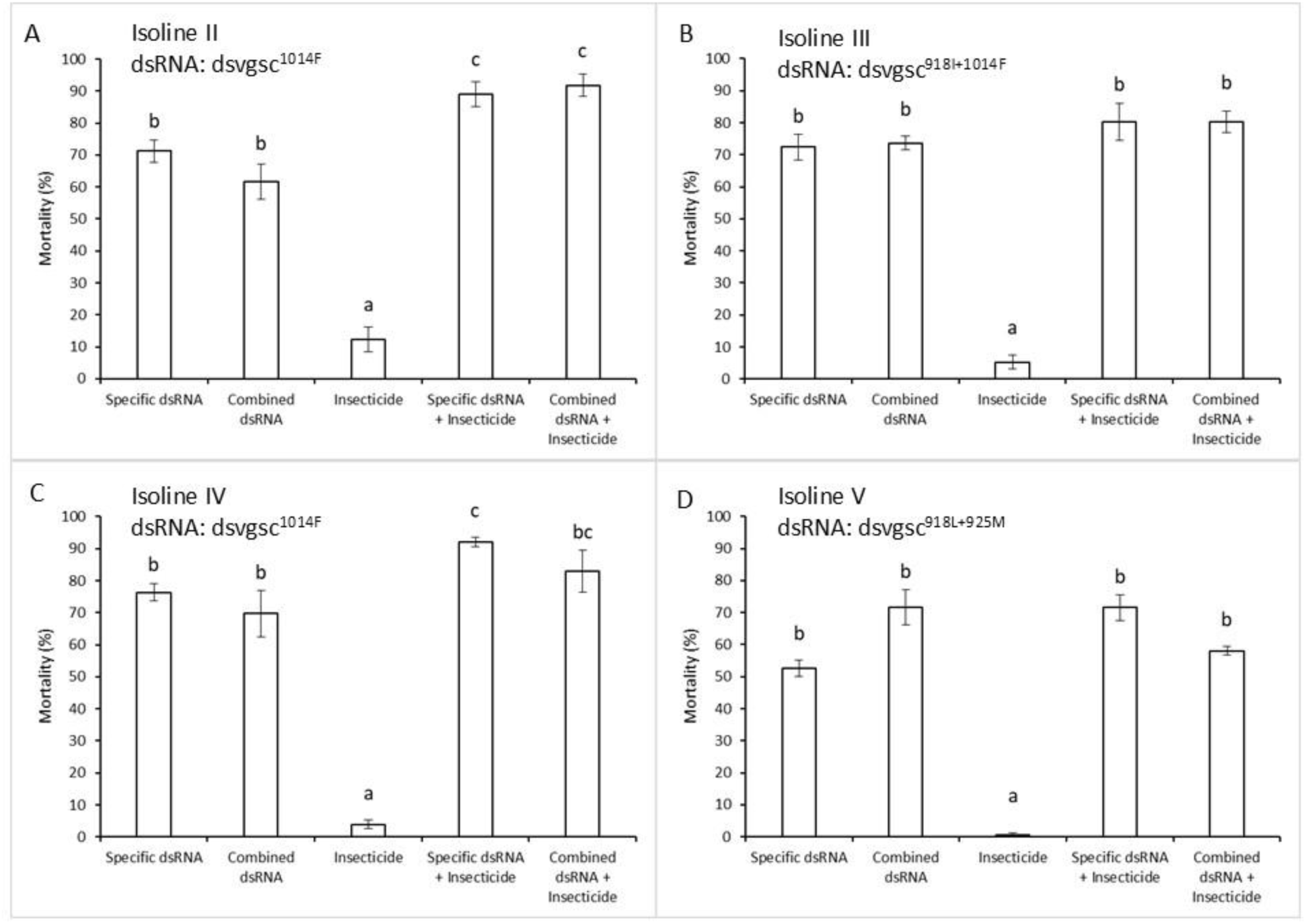
Mortality induced in pyrethroid-resistant *Aphis glycines* isolines II, III, IV, and V (Table 1) when exposed to a constant 1.0 µg µl^-1^ concentration of a specific dsRNA whose sequence matched the resistance alleles in the corresponding isoline (Specific dsRNA; Table 2) or a dsRNA combining all nucleotide substitutions across all resistance alleles (Combined dsRNA; Figure 1). Specific and Combined dsRNAs were applied alone (dsRNA) or along with lambda-cyhalothrin at a constant concentration of 1.23 ppm (dsRNA + Insecticide). Mortality was corrected using Abbott’s formula based on survivorship in untreated controls. Different letters among treatments within isoline indicate significant differences between pairwise comparisons according to Šidák tests.

The greatest amount of mortality observed when only the insecticide was used occurred with aphids from a *w*+ susceptible Isoline (i.e. Isoline I). When Isoline I was treated with 1.23 ppm lambda-cyhalothrin, 92% mortality was observed (Figure 4A). The treatment of Isoline I with Combined dsRNA alone resulted in 55% mortality (Figure 4A), which was significantly lower than when Isoline I was treated with the insecticide alone. Exposure of aphids from Isoline I to the Combined dsRNA with the insecticide did not significantly increase the mortality compared to the insecticide alone. The production of nymphs from Isoline I was affected by exposure to the insecticide alone (Figure 4B). When Isoline I was treated with the Combined dsRNA, nymphal production was numerically intermediate but statistically indistinguishable from either the untreated control or the treatments including insecticide.

**Figure 4.**
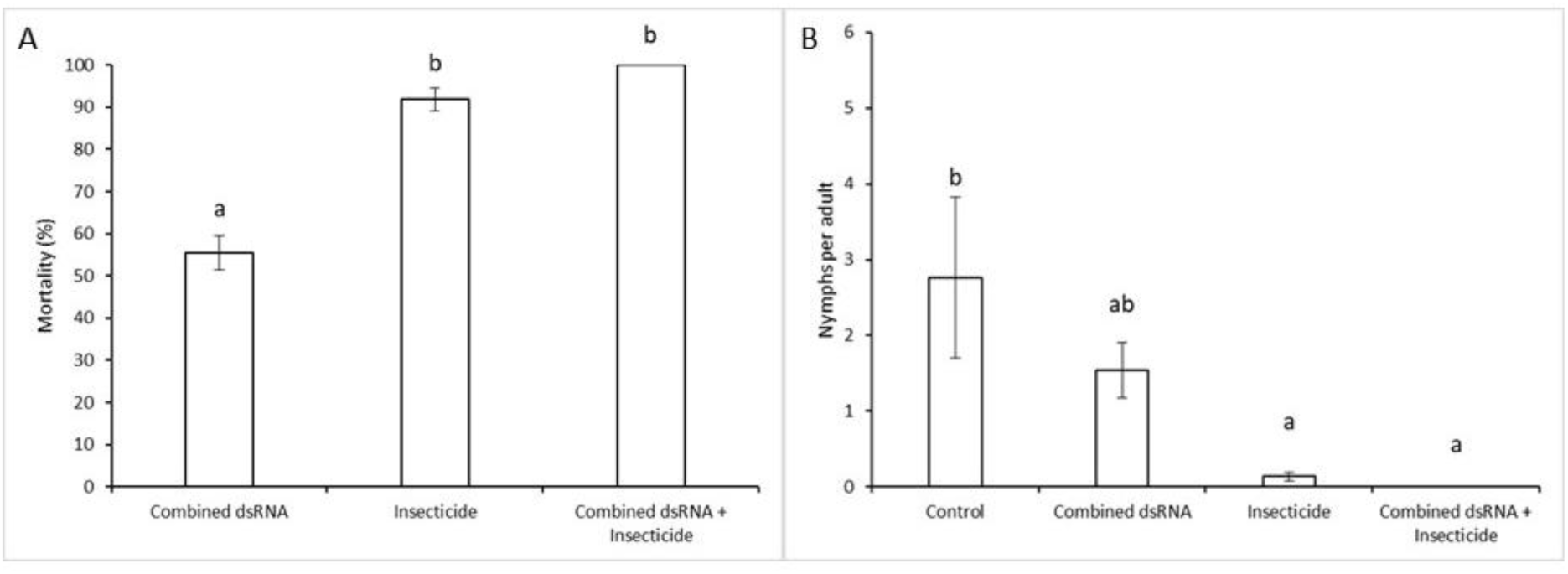
Mortality (A) and nymphs produced (B) in the pyrethroid susceptible *Aphis glycines* Isoline I (Table 1) when exposed to a double-stranded RNA (dsRNA) that combined all nucleotide substitutions across all resistance alleles (Combined dsRNA; Figure 1). Each dsRNA treatment was conducted both in the presence (+ Insecticide) and absence of 1.23 ppm of lambda-cyhalothrin. Mortality was corrected using Abbott’s formula based on survivorship in untreated controls, and nymphs per adult were calculated as the total number of nymphs divided by the number of adults alive 24 h after treatment. Different letters indicate significant differences according to Šidák tests.

We observed significant variation in nymph production among the resistant Isolines when exposed to dsRNA, both alone and in combination with insecticide (Figure 5). The greatest number of nymphs produced was observed among the controls, across isolines II, III and V and among aphids from Isoline IV treated with the insecticide alone. Except for Isoline II, nymphs production did not vary among treatments with insecticide alone compared to the controls (Figure 5A). Treatments with Specific and Combined dsRNAs alone and with the insecticide demonstrated varying effects on the number of nymphs per adult. Nymph production was significantly lower from isolines III and V when treated with their Specific and Combined dsRNA alone and with their Specific dsRNA with the insecticide compared to control or insecticide treated counterparts (Figure 5B, 5D). Nymphs produced by Isoline III or IV females treated with the Combined dsRNA with the insecticide were not significantly reduced compared to treatments with the insecticide alone. Reproduction was also not significantly reduced among females from Isoline II or IV when exposed to their Specific dsRNA or Combined dsRNAs alone (Figure 5A, 5C). Treatment of Isoline II females with Specific and Combined dsRNA with the insecticide reduced nymphal production to nearly zero (Figure 5A). Among females from Isoline IV, only those exposed to the Combined dsRNA alone or the Specific dsRNA + Insecticide showed a significant reduction in nymph production compared to treatment with insecticide alone. In isoline IV, the Specific dsRNA with insecticide treatment resulted in the lowest reproductive rate.

**Figure 5.**
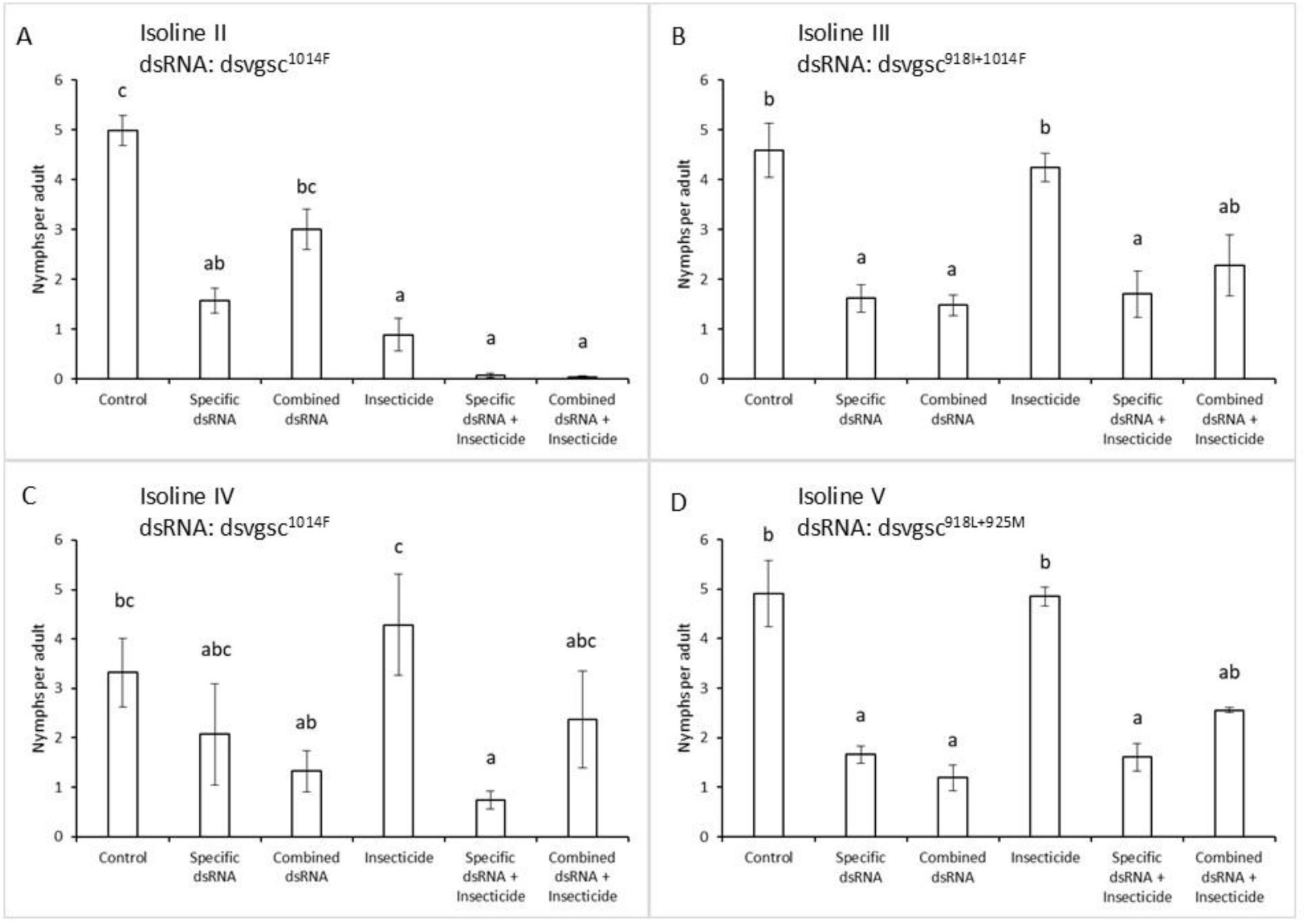
Number of nymphs produced per adult across pyrethroid resistant *Aphis glycines* Isolines II, III, IV and V (Table 1) when exposed to a dsRNA matching the resistance allele in their corresponding genotype (Specific dsRNA; Table 2) and to a dsRNA that combined all nucleotide substitutions across all resistance alleles (Combined dsRNA; Figure 1), and when both dsRNA treatments were applied alone or in combination with the insecticide lambda-cyhalothrin at a constant 1.23 ppm concentration (dsRNA + Insecticide). Nymphs per adult were calculated as the total number of nymphs divided by the number of adults alive 24 h after treatment. Different letters among treatments within an isoline indicate significant differences according to Šidák tests.

## 4 DISCUSSION

The current study demonstrates that dsRNAs targeting specific alleles of the *vgsc-h1* gene associated with pyrethroid resistance in *A. glycines* cause mortality in a dose-dependent manner. Specific dsRNAs alone cause significantly greater mortality among aphids in isolines carrying the corresponding resistance allele when compared to treatment with insecticide alone. This restoration of susceptibility across lines agrees with our prior findings that validate the role of *vgsc-h1* in the pyrethroid resistant phenotypes,^37^ a role that is supported by genetic associations among insects^12^ and several aphid species,^15–19^ including *A. glycines*.^14,20,21^ The Specific dsRNAs acted synergistically with a pyrethroid (lambda-cyhalothrin) to statistically increase mortality in two of the four aphid isoline-Specific dsRNA pairings tested (Isolines II and IV; Figure 3). The lack of statistical significance for Isolines III and V is likely due to a high level of mortality caused by the dsRNAs alone. RNAi-mediated *vgsc* transcript knockdown was previously shown to increase pyrethroid susceptibility in a resistant strain of the mosquito *Aedes aegypti* (Diptera: Culicidae), functionally validating the role of this gene in phenotypic resistance and proposing exploitation as an insecticidal RNA.^43^ Insecticidal properties of dsRNAs targeting *vgsc* orthologs were also previously realized through the dose-dependent survival probabilities of up to nearly 50% among larvae of the red flour beetle, *Tribolium castaneum* (Coleoptera: Tenebrionidae) at 2 days post injection or oral ingestion.^44^ Our treatments with allele-specific dsRNA led to similar percentage mortalities (≥ 52%) at 2 days post topical application (Figure 3), but direct comparison between experiments and within a study is challenging due to a multitude of factors that influence efficacy including differences in application method, variation in dsRNA uptake by different species,^30,45^ varying properties of RNA sequences that influence processing by the RNA-induced silencing complex (RISC),^46^ and sequence match to transcript variants in a population.^47,48^

Despite these complexities, we investigated the specificity of dsRNA targeting *vgsc-h1* allelic variants among the four distinct genotypes in our pyrethroid resistant isolines. Our experiments show no significant effects of sequence mismatch. This is revealed by comparing mortality induced in each resistant isoline by the Specific dsRNA, (i.e. 100% nucleotide sequence identity to the resistant alleles), and by the Combined dsRNA (ds*vgsc-h1*^All^), with a consensus sequence of all mutant alleles (Figure 3). Our result might be surprising considering differences in zygosity (copy number) of the corresponding resistance allele and degree of sequence mismatch among the isolines, but not when variance in estimated number of mismatches between isolines is considered. Specifically, alleles conferring pyrethroid resistance in *A. glycines* are dominant, where heterozygotes in Isolines III and V carrying one copy of the *w*+ allele can show phenotypes (RR) similar to the homozygote Isoline IV (Table 1).^14^ Provided that the *w*+ allele in each heterozygote is predicted to produce four mismatches with the Combined dsRNA and between two and three mismatches with any of the Specific dsRNA, the total difference in number of mismatches among strains varies by only two of the total eight possible (25%) across both alleles. Provided that ds*vgsc-h1*^All^ is 485 nt in length, the six to eight total mismatches result in 1.2 and 1.6% sequence differences. This partially agrees with prior findings that showed mortality induced by a 200 nucleotide long dsRNA was not significantly reduced even when six mismatches were introduced.^48^ Analyses of introduced mismatches demonstrate that RNAi-mediated knockdown can be effectively triggered by dsRNAs with at least 80% sequence identity to the target.^49^ Thresholds of ≥16 nucleotide stretches of 100% identity or ≥26 regions with over 93% identity with ≥ 5 nucleotides between mismatches can trigger an RNAi response. These parameters dictating potential efficacy suggest that the number and spatial arrangement of nucleotide mismatches between the Combined dsRNA and *w*+ or resistant alleles are unlikely to impact knockdown and subsequent mortality. This further suggests that any of the dsRNAs used in this study (Table 2) might lead to similar mortalities across *vgsc-h1* genotypes found in field populations.^14,20,21^

We also observed significant impact of dsRNA on *A. glycines* reproduction. Females reproduce asexually through parthenogenesis during spring and summer when infesting *G. max*, giving live birth to clonal daughters. This can result in high population growth rates under ideal conditions,^11,50^ producing outbreaks that cause significant yield loss.^1,4,5^ We observed that both Specific dsRNAs and the Combined dsRNA applied alone and with lambda-cyhalothrin significantly reduced the number of nymphs per treated pyrethroid resistant female (Figure 5). This phenomenon was most pronounced in Isolines III (Figure 5B) and V (Figure 5D), where in both the Specific dsRNAs and the Combined dsRNA applied alone, and the Specific dsRNAs with lambda-cyhalothrin significantly reduced nymph production compared to the control and insecticide treatments. A significant reduction in nymphs of Isoline IV was also observed when aphids were treated with Specific dsRNAs and insecticide compared to the insecticide alone (Figure 5C). These comparisons collectively demonstrate the impact that our dsRNAs have on reproduction of pyrethroid resistant aphids when applied with lambda-cyhalothrin, but not observed by exposures to the insecticide alone. This may not be surprising given the systemic effects of RNAi and telescoping generations of aphids contained within a single female aphid. Specifically, developing daughters within treated females are likely to absorb the dsRNA, creating a combination of negative impacts on the vigor of the mother and her developing daughters up to and after birth. Treated females in this study were not dissected to detect moribund progeny, nor were reproductive effects on individual surviving daughters measured. Regardless, evidence suggests there may be multigenerational effects on aphid reproduction, and future studies are needed to further characterize their impacts. Similar negative reproductive effects were detected in the aphid *Myzus persicae* that fed on *Nicotiana benthamiana* that produced dsRNA targeting the aphid’s *rack-1* and *MpC002* genes,^51^ and among females of *A. gossypii* following dsRNA-mediated knockdown of the sucrase gene.^52^ These lines of evidence suggest that the effects of insecticidal dsRNA treatments may extend to clonal daughters and function to further suppress pest aphid populations.

Pyrethroid resistance has evolved via several mechanisms independent of target site mutations, such as behavioral avoidance, reduced cuticular penetration, and increased detoxification.^53^ Indeed, a prior study associated the constitutive or induced up-regulation of 11 transcripts from putative detoxification genes (esterases, carboxylates and cytochrome P450 monooxygenases) with pyrethroid resistant field populations of *A. glycines* in southern Minnesota.^54^ The up-regulation of transcripts from putative detoxification genes was also shown in a laboratory strain of *A. glycines* collected from fields in northern China with a 76.67-fold increase in lambda-cyhalothrin resistance.^55^ The role of detoxification as the resistance mechanism of this laboratory strain was validated by restoration of susceptibility equivalent to that of a control strain when chemical esterase and cytochrome P450 inhibitors were applied along with lambda–cyhalothrin. These prior lines of evidence suggest that genotypes underlying these traits could be heterogeneous, realized through the contribution of different mutations at one or more genetic loci. In contrast, chemical inhibitors of cytochrome P450, esterase, and glutathione-S-transferase activities all failed to increase pyrethroid susceptibility among resistant *A. glycines* in the isofemale line SBA-MN1-2017-ISO (Isoline III; Table 1). Susceptibility to lambda-cyhalothrin increased significantly in Isoline II when treated with dsRNAs targeting *vgsc-h1* transcripts (Figure 3), validating the role of *vgsc* mutations in the phenotypic resistance observed in this isoline. Differences among these studies and our current results could reside in sources of phenotypic resistance; a set of distinct homogenous genotypes in isofemale lines with field-evolved resistance compared to a heterogeneous pool of genotypes subjected to incremental increases in dose under an artificial selection regime. Laboratory selection can lead to fixation of multiple genes, each with small additive or non-additive effects on the phenotype, whereas strong directional selection in outcrossing field populations tends to arrive upon a single gene with a major effect.^56^ This phenomenon, along with validation of phenotypic effects of ds*vgsc-h1* knockdown on isolines with field-derived resistance shown in this study, suggests that non-synonymous mutations in the D2S4-6 pyrethroid target site of the *A. glycines vgsc-h1* gene constitute a major contribution to explaining insecticide-resistance in field populations.

Foliar applications of pyrethroids are the most widely used tactic to manage *A. glycines* populations in the Midwest United States,^17,57^ where comparatively low cost of pyrethroids bolsters profit margins for farmers.^5^ Tactics preserving the efficacy of pyrethroids in geographic regions where resistance pervades could thereby extend the associated economic benefits. In this study, we demonstrate that topically applied dsRNAs targeting the *A. glycines vgsc-h1* not only lead to greater mortality among pyrethroid resistant aphids compared to treatments with lambda-cyhalothrin but can function synergistically with lambda-cyhalothrin to cause further increase in mortality. Technologies that synergize with insecticidal agents compromised by resistance pest populations can restore efficacy, offering tools for insect resistance management.^58^ Restoring pyrethroid efficacy in resistant field populations remains a relatively novel concept, where one prior implication of such a tactic has been found.^43^ Mortality caused by our dsRNAs applied in the laboratory was accompanied by the unexpected effect of reducing the number of nymphs produced by females, suggesting that beyond initial mortality, these putative longer-term effects on reproduction may function to further suppress population growth. Future investigations into these suppressive effects we demonstrated in laboratory conditions are needed to confirm their impact under field conditions.

## ACKNOWLEDGEMENTS

This work was supported by a grant from the Iowa Soybean Association Award, “Advancing insect pest management for soybeans: Combating the soybean gall midge and restoring susceptibility to insecticide resistant insects”. This research was completed with support from the USDA, Agricultural Research Service (ARS) CRIS Project 5030-30400-019-000D. Mention of trade names or commercial products in this publication is solely for the purpose of providing specific information and does not imply recommendation or endorsement by the USDA. USDA-ARS is an equal opportunity employer and provider. The dsRNA sequences reported here are part of the application PCT/US24/59991, Compositions and methods for restoring susceptibility to pyrethroid insecticides in resistant populations, filed at the United States Patent and Trademark Office on 13 December 2024.

## Notes

### Competing Interest Statement

The authors have declared no competing interest.

