## Supplementary material for "Reversion of pyrethroid resistant phenotypes in *Aphis glycines* by topical delivery of dsRNA targeting resistance alleles at the vgsc locus": Suplementary material

### Supplementary materials

Table S1:

```
A)
AG6007485_2711t7-S   TAATACGACTCACTATAGGGAGAGTTCATTTCGTTTGCTTCGAGT
AG6007485_3195t7-A   TAATACGACTCACTATAGGGAGAGAATCGGCCAATACGTTCAAA

B)
Wt susceptible
GTTCATTTCGTTTGCTTCGAGTATTTAAGTTGGCAAAATCTTGGCCCACACTTAATCTTTTAATATCAATAATGGGTCGAACCATTGGTGCTTTGGGTA
AgΔ918I – Boone template
GTTCATTTCGTTTGCTTCGAGTATTTAAGTTGGCAAAATCTTGGCCCACACTTAATCTTTTAATATCAATAATAGGTCGAAC
AgΔ918L – Boone template                                     SspI (AAT^ATT)
GTTCATTTCGTTTGCTTCGAGTATTTAAGTTGGCAAAATCTTGGCCCACACTTAATCTTTTAATATCAATATTGGGTCGAA
AgΔ925M – Boone template
GTTCATTTCGTTTGCTTCGAGTATTTAAGTTGGCAAAATCTTGGCCCACACTTAATCTTTTAATATCAATAATGGGTCGAACCATTGGTGCTATGGGTA
AgΔ918+925 Use Kanawha 1014F homozygote cDNA as starting.
GTTCATTTCGTTTGCTTCGAGTATTTAAGTTGGCAAAATCTTGGCCCACACTTAATCTTTTAATATCAATATTAGGTCGAACCATTGGTGCTATGGGTA
```

**Need to generate:**
**Homozygous MN1 918L + 1014F probe**

```
AgΔ918I – Kanawha template
GTTCATTTCGTTTGCTTCGAGTATTTAAGTTGGCAAAATCTTGGCCCACACTTAATCTTTTAATATCAATAATAGGTCGAAC
```

**Homozygous Darwin 918L + 925M probe**

```
AgΔ925M – Darwin primer extension #1 as template
GTTCATTTCGTTTGCTTCGAGTATTTAAGTTGGCAAAATCTTGGCCCACACTTAATCTTTTAATATCAATAATGGGTCGAACCATTGGTGCTATGGGTA
GTTCATTTCGTTTGCTTCGAGTATTTAAGTTGGCAAAATCTTGGCCCACACTTAATCTTTTAATATCAATATTGGGTCGAACCATTGGTGCTTTGGGTA.... Template a
GTTCATTTCGTTTGCTTCGAGTATTTAAGTTGGCAAAATCTTGGCCCACACTTAATCTTTTAATATCAATAATGGGTCGAACCATTGGTGCTATGGGTA.... Template A
Then
AgΔ918L – Darwin template primer extension #1                SspI (AAT^ATT)
GTTCATTTCGTTTGCTTCGAGTATTTAAGTTGGCAAAATCTTGGCCCACACTTAATCTTTTAATATCAATATTGGGTCGAA
GTTCATTTCGTTTGCTTCGAGTATTTAAGTTGGCAAAATCTTGGCCCACACTTAATCTTTTAATATCAATATTGGGTCGAACCATTGGTGCTATGGGTA...
GTTCATTTCGTTTGCTTCGAGTATTTAAGTTGGCAAAATCTTGGCCCACACTTAATCTTTTAATATCAATAATGGGTCGAACCATTGGTGCTATGGGTA...
```

Figure S1. Transcript model AG6007485-RA annotated in the genome assembly for *Aphis glycines* biotype 1 (Ag_bt1_v6.0; Giordano et al. 2020) putatively encoding the wildtype (*w*+) allele of the *voltage-gated sodium channel* subunit h1 (*dsvgsc-h1*) gene (Valmorbida et al. 2022a). Start (ATG) and stop codons (TAG) are highlighted. Nucleotides from positions 2633 to 3195 are encompassed within the *dsvgsc-h1*$^{w+}$ double-stranded RNA (dsRNA; **Figure 1**), and nucleotides that are changed among native alleles associated with pyrethroid resistance in *A. glycines* populations and among allele-specific dsRNA are in bold.

*Aphis glycines vgsc-h1* CDS from gene model AG6007485-RA; pyrethroid susceptible allele.

ATGTCCATTGCTGACACCGATTCTTCATTCTCCAGTGAAGAAAAAAGCCTATTCCGTCCGTTCACCCGCGAATCACT
GAGGGCCATCGAACAACGCATAGCTGAAGATCATGCTAAACAGAAAGAACTTGAGGAAAAAAGAGCTGAAGGGGAGC
CCTTGCCATGGAAGAGAAATAAAAATAAAGACGTTCGGTATGAGGACGAGGACGAAGATGAAGGTCCCCAGCCGGAC
GCTACTCTGGAACAAGGAGCTCCGCTGCCGGTCCGTTTGGTCGGCACGTTTCCACCGGAATTAGCGTCCGTACCGCT
CGAGGATATCGACCCTTACTATCACAATCAAAAAACTTTTGTAGTAATTAGCAAAGGAAAAGATATCTTCAGGTTCA
GCGCCACCGATGGCCTCTGGGCACTCGACCCCTTCAACCCTATCCGCCGGGTGGCTATTTACATATTAGTCCACCCG
ATATTTTCTGTAACAATCATCACTACTATACTTACAAACTGTGTGTTCATGATAATGCCCCCAACTCCGACTATTGA
AGCGTCTGAAGTAATATTTACCGGCATCTACACATTCGAATCGGCTGTGAAAGTAATGGCCCGAGGTTTCATATTAG
AACACTTCACCTATCTTAGAGATGCATGGAATTGGCTAGACTTCATTGTTATTGCATTAGCTTACGTTACTATGGGT
ATAGAACTTGGAAATTTAGCGGTTCTTCGAACATTTCGAGTACTGCGAGCGCTCAAAACTGTAGCCATTGTGCCTGG
ATTAAAGACTATCGTTGGAGCTGTGATAGAATCCGTGAAAAACCTCAGGGATGTCATTATATTAACAATATTTTCAC
TATCTGTGTTCGCATTACTGGGATTACAAATTTATATGGGCGTATTAACACAAAAATGTATAAAATATTTTCCTCTT
GACGGCTCAGCTGGAAATTTAACCAATGAAAATTGGTTTGCGTTCATGTCGAATAAATCTAACTGGCAACCTAATGA
CGATGAACCAGATGAGTATCGATTATGTGGGAACGGTACTGGCGCTGGACAATGCGATGAAGGTTATATGTGTATCC
AAGGCTTTGGAGGAAATCCTAACTATGGATACACAAGTTTTGATACATTTGCGTGGGCTTTACTATCGGCATTTAGA
TTAATGACTCAAGACAACTGGGAAGCATTATACCAACAGGTGTTAAGAGCTGCCGGACCATGGCATATGTTCTTCTT
CATTGTGATTATATTTCTCGGTTCGTTTTATCTTGTCAATTTGATATTAGCCATCGTAGCAATGTCGTACGACGAAT
TGCAGAAAAAAGCTGAAGAAGAAGAGGCAGCCGAAGAAGAAGCTATCAGGGAAGCGGAACAAGCGGCTAAAGACAGA
GAGGTCCGGAGGCAAGCTCACGAGGAAAGAGTGGCGGAACGTGCGGAGAGAGCGCGACACGTCACCCAACATCCGAA
ATCGCCTTCGGACTTTTCGAGCCAAAGTTACGACAATATGTTTGGCGGTGGTCAAGACCGAGGGATTGGCAATGATC
ATCATAGAGAGAAAAATATGAGCCTGCGATCGATGTCGATCACGAGCCATGACAAACATTCGGACACCGGGAGTGTG
GATAGGCAGAGCGGCAAAACAAGAAAAGCAAGCCTCAGCCTGCCGGGATCGCCGTTCAACATTCGGCGGGCGTCGCG
TGGCAGTCACCAACTGTCACATCGGAACGGTAGACCGAGGTTTACCGGAGCGGATACCAAGCCGTTAGTCTTAAACA
CGTTACTTGACGCGGAAGAACATTTACCGTATGCCGACGACTCAAACGCAGTGACGCCCATGAGTGAGGAGAACGGT
GCCATCATCGTACCCGTATCATACGCGAACTTCGGTTCGAGGCACAGCAGTTACACGTCACACACGTCCCGAATTAC
GTATACGTCGCATGCGGATCTGTTCAAACCGCCTATGACCAAAGAGAGGCAGTTGAGGTCGCGGTCGGCCAGAAACT
ATTTCAATCCATCTGATCAGAGATACCATCGAGACGACGATTACGATACGAGTTCCATGTCCAAATCGAAACAAAAA
GTGGACGAGTGCGGTTACAGTGATTCACAAAAGCACACGGTCGTCGACATGAGAGACGTTATGGTGTTGAACGATAT
CATCGAACAGGCAGCCGGACGTCAGAGTAGGGGTAGCGAAAAAGCAGTAATAACGGGGGGTACTCTCGTGGGTGCAT
GGGGTACAGTGTCCACGGTGTACGTGTTCCCAACAGATGAAGATGCGGTCGACGGAGAAGACGAAGAGGAAGAAAAT
GAGGAACCCACTTTTCGTGAAAAGTTCCAAGTATGGTTATTGAAGTTCATCGACACGTTTTGTGTCTGGGACTGTGG
ATGGCCGTGGCTCAAGTTTCAGCAAGGACTCGCATTCATAGTGTTTGATCCATTCGTCGAACTCTACATCACCCTGT
GTATTGTAGTTAACACGTTATTTATGGCGCTCGATCATCATGAAATGGATCCCAAATTGGATTTCATACTCAACAAG
GCTAATGTTTTTTTCAGTGCTACGTTCGGCGTTGAAGCAGCTCTGAAACTTATGGCTATGAGTCCTAAGTATTACTT
CCAAATGGGCTGGAACATCTTTGACTTCATCATCGTAATTCTTTCCGTAGTAGAATTGCTCTCCGCGGGTTACCAAG
GACTTTCCGTATTGCGTTCATTTCGTTTGCTTCGAGTATTTAAGTTGGCAAAATCTTGGCCCACACTTAATCTTTTA
ATATCAATA**A**T**G**GGTCGAACCATTGGTGCT**T**TGGGTAACCTAACGTTTGTGTTGTGCATAATCATATTTATATTCGC
CGTTATGGGTATGCAGTTATTTGGAAAAAACTACACAGAAAAAATGTACTTATTCAAAGACCACGAGCTTCCCCGGT
GGAACTTCACCGATTTTTTGCACTCGTTTATGATAGTATTTCGAGTATTATGTGGTGAATGGATTGAATCAATGTGG
GACTGCTTACACGTTGGAGAACCAACGTGTATACCATTCTTCTTGGCTACTGTTGTCATCGGTAAC**C**TTGTGGTACT
TAATCTTTTCTTGGCGTTGTTGCTGAGTAATTTTGGCTCGTCTAATTTATCGGTGCCTACGGCTGATAGCGACACAA
ACAAGATCACAGAAGCATTTGAACGTATTGGCCGATTCAATAAATGGGTGAAAACGCATATTCTGAACTTCCTCAAA
GTACTCCGACTGAAAATCACCAATCAAATATCGGTTCAAGTGTCAGGTCGTGACAGAGATATTGACTTGCCTGTTGA
CGAAACAATTGTAGATGTTATTGCGCCATTTAAAGACACTAAGGAACCAGTAGAGATGACAATTGGTGACGGGATGG
AATTCACGATTCCGGGTGATGTAAAACAAAAAATAAAAAAGAACCAAGTTGGGAATTCAATTGGAAATCATCAAGGG
AACAAAGTTGGAAACGATTACAAAAAGGAAAGTTTCGACCTGGATAGTTTAAAATGCTAG
